# Immune-neuroendocrine phenotypic expression changes through life in *Coturnix japonica* quails

**DOI:** 10.1101/2024.01.24.577005

**Authors:** Antonela Marozzi, Silvia G. Correa, Rupert Palme, Veronica I. Cantarelli, Marina F. Ponzio, Raul H. Marin, F. Nicolas Nazar

**Author notes:** Corresponding Authors: R. H. Marin; F. N. Nazar. Raul Marin and F. Nicolás Nazar should be considered joint senior author.

## Abstract

Immune-neuroendocrine characteristics can be used to classify individuals according to their physiological profiles or phenotypes (INPs). In avian models such as quail and domestic chickens, three subgroups based on INPs have been defined: Lewis-like (pro-inflammatory polarization), Fischer-like (anti-inflammatory polarization), and an intermediate INP. This study investigates the stability and alterations of INPs throughout ontogeny, from juvenile to adult stages in four time-points including an exposure to unpredictable and diverse chronic stress (CS) during early adulthood. We measured corticosterone levels, pro-(IFN-γ and IL-1β) and anti-inflammatory (IL-13, IL-4) cytokines, phytohemagglutinin (PHA-P) lymphoproliferative response, anti-sheep red blood cells antibody (Ab SRBC) response, and leukocyte distribution frequency. Cluster analyses were conducted to classify bird based on their similarities across all analyzed variables, to thereby establish their INP at each time point. The extreme Lewis- or Fischer-like profiles were less represented in juvenile and pre-stress adult birds showing a higher proportion of individuals with an intermediate profile. Following CS exposure, the prevalence of Lewis-like and Fischer-like profiles increased. This shift persisted 10 weeks later as birds matured to an advanced egg-laying stage, with females predominantly exhibiting the Fischer-like INP, and males the Lewis-like INP. The observed shift in INP distribution following CS towards more polarized Lewis- and Fisher-like profiles implies a more even representation of the three observed profiles and may reflect inter-individual differences in physiological response to CS associated to particular coping strategies. A more even INPs distribution could provide the population with a greater advantage when facing diverse environmental challenges.

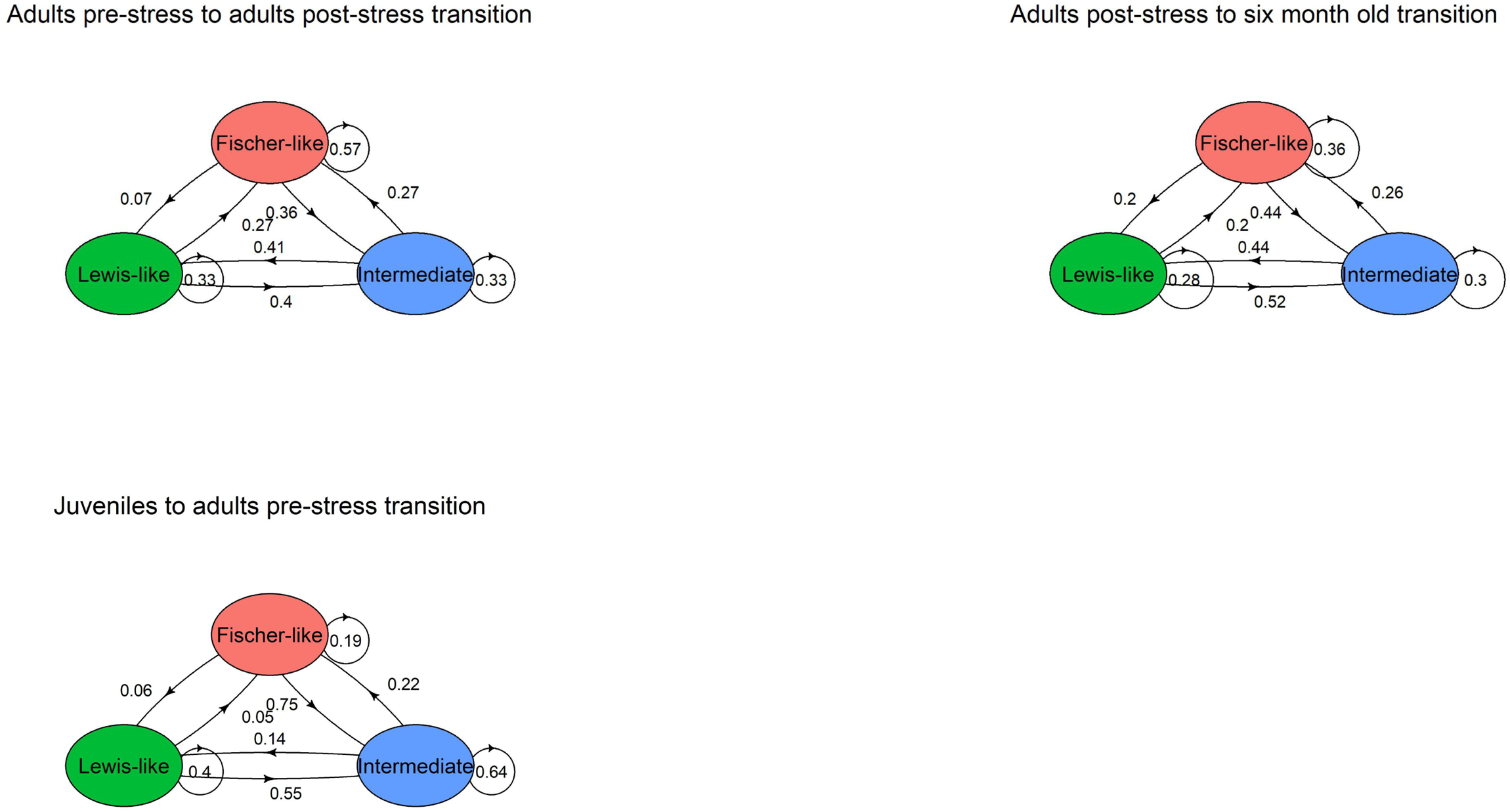

## Introduction

Populations are variable, and this variability is critical to coping with internal demands (e.g., growth and development) and external demands (e.g., environmental changes) (Giayetto et al., 2020; Nazar et al., 2017; Randolph et al., 2024; Reed et al., 2010). Consequently, the interaction of these factors may shape the physiological configuration of any population (Martin, 2009). In general, when juveniles face environmental changes such as variations in average annual temperature, food availability or handling, these can lead to shifts in their growth, weight gain, potentially causing permanent changes in their phenotype configuration (Piersma and Drent, 2003; Stearns, 1989). In contrast, adults tend to develop what is known as a “reversible state effect” where they can switch between phenotypes, adopting new configurations in response to environmental changes (Burggren, 2018; Senner et al., 2015). To differentiate between the permanent changes in reaction norms that occur during juvenile stages and the reversible adaptations that happen in adulthood, Piersma and Drent (2003) proposed the term “phenotypic flexibility”. This concept refers to changes that occur when environmental conditions vary rapidly and over shorter timescales than a lifetime. The ability to adapt by acquiring a reversible state effect can be an evolutionary advantage, as greater trait diversity within a population enhances its preparedness for future unknown challenges (Forsman, 2015).

Modifying phenotype configuration can lead to a reorganization of resource allocation, which may impact fitness (Buehler et al., 2010; Romero et al., 2009). In some cases, these effects are immediate, while in others, they may lead to carry-over effects that influence an individual’s performance in future situations (O’Connor et al., 2014), potentially affecting population-level processes such as survival rate (Loonstra et al., 2019; Saino et al., 2017). Even so, in some cases, carry-over effects are not seen until the next generation, where the price of facing a challenge becomes evident in the descendants (Giayetto et al., 2020; Senner et al., 2015).

Immune neuroendocrine phenotypes (INPs) can be used to understand at least part of the physiological variability. They were described in mammalians including humans (Elenkov et al., 2008), and related species like rodents (Wei et al., 2003). Later on, they were described in avian species, such as *Gallus gallus* (Nazar *et al.*, 2017), and *Coturnix japonica* (Nazar, 2015). INPs represent a set of characteristics that define individuaĺs neuro-hormonal and immunological responses. These responses are within the same phenotype but differ from those of other phenotypes within the same population (Sternberg *et al.*, 1989; Elenkov *et al.*, 2008; Ashley and Demas, 2017). Consequently, INPs provide a valuable framework for assessing avian populationś responses to chronic and stressful environmental challenges (Sternberg *et al.*, 1989; Nazar *et al.*, 2017). INPs are characterized by specific concentrations of neuroendocrine mediators, activity and expression of hormonal receptors, and cytokine profiles that are either pro- or anti-inflammatory. In quail and hens, three INPs have been described: (i) the Lewis-like phenotype, characterized by low corticosterone levels, high levels of inflammatory mediators (IFN-γ and IL-1β), a strong lymphoproliferative response to Phytohemagglutinin-p (PHA-P), high antibody response against to sheep red blood cells (SRBC), a low innate-to-acquired immunity ratio as evaluated by the frequency of leukocyte distribution (FLD), and anti-inflammatory mediators (IL-13, IL-4); (ii) the Fischer-like phenotype, characterized by high corticosterone levels, high levels of anti-inflammatory mediators (IL-4 and IL-13), high LDF and low lymphoproliferative response to PHA-P, Ab SRBC, and proinflammatory response (Nazar et al. 2015); and (iii) an the intermediate phenotype that lies between these two extremes (Fig. 1). INPs were mainly described in a comparative context, e.g., by comparing groups of individuals with high basal levels of inflammatory cytokines to others with significantly lower levels within the same species. Identifying different INPs within a population is crucial, as they can influence the ability of individuals to cope with environmental challenges i.e., from stress associated with maintenance chores to disease (Ashley and Demas, 2017; Elenkov et al., 2008; Nazar et al., 2015, 2017). Specifically, the prevalence of specific INPs within a population can influence how well individuals with those phenotypes respond to immune neuroendocrine (INE) challenges, such as the presence of pathogens (Nazar et al. 2017; Nystrand and Dowling 2020). Additionally, INPs reflect underlying differential INE matrix response, i.e., activity in the hypothalamus-pituitary-adrenal axis (HPA) or in the sympathetic-adrenergic axis (Elenkov et al., 2008, 2000; Nazar et al., 2015, 2017), and may consequently influence on metabolic rate (Jimeno et al., 2018b) and behavior (Bókony et al., 2009).

**Fig. 1:**
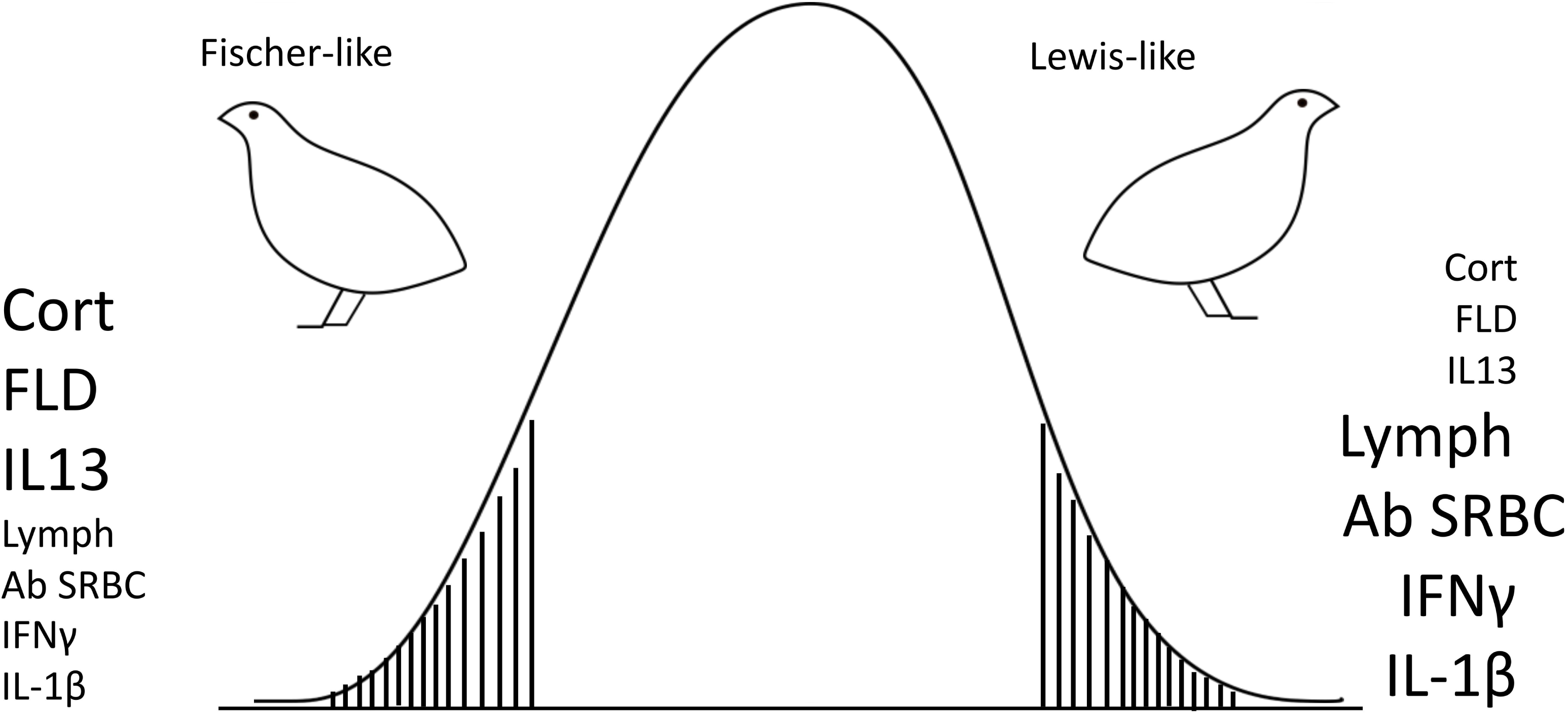
Conceptual schematic representation of INPs in *Coturnix japonica* (adapted from Nazar et al. 2015). The variables used to define avian INPs are displayed beside each bird. The size of each variable indicates whether the animal exhibits high or low responses in these parameters. LYMPH: lymphoproliferative response to PHA-P; Ab SRBC: antibody response against SRBC; FLD: frequency of leukocyte distribution; level of expression of mediators: IFN-γ and IL-1β (pro-inflammatory); and IL-4 and 13 (anti-inflammatory). “Fischer-like” quail with high CORT levels also manifest high FLD and IL-13, but low Lymph, Ab SRBC, IFN-γ and IL-1β levels. “Lewis-like” counterparts have low CORT as well as low FLD and IL-13 responses, but high Lymph, Ab SRBC, IFN-γ and IL-1β responses. Notably, both groups do not differ in their IL-4 levels.

Also, the effect of sex may be an important driver on INE response due to the effect of estrogens and testosterone (Garcia-Reyero, 2018; Klein and Flanagan, 2016). Females tend to allocate more resources to reproduction than males, given the demands of egg development and parental care (Lee, 2006). As a result, some studies support the idea that females tend to exhibit a Th2-biased immune response, i.e., potentiated humoral response less expensive in terms of energy budget and more related to extracellular parasites such as helminths (Lee, 2006; Vincze et al., 2022). On the other hand, males tend to invest more energy in mating attraction, which some studies suggest allows them to allocate more energy to Th1-biased immunological response, characterized by cell-mediated immunity (Lee, 2006; Valdebenito et al., 2021). Th1 cells are particularly effective in protecting against intracellular infections by viruses and bacteria (Kogut, 2022). In the particular avian model used in this study, two key considerations emerge: first, these animals have undergone intense selection for egg or meat production, which may partly influence their INE responses; second, certain physiological responses, critical for survival, may remain resilient to reproductive selection pressures, persisting even in highly selected populations (Ashley and Demas, 2017; Cheng, 2010).

In summary, INPs expression could be variable through life, giving more flexibility to the population to respond to environmental variability (Senner et al., 2015; Stager et al., 2024). Based on the previously described intrinsic population variability and the individuals’ capacity to alter phenotypic expression, we investigated the variability in INP expression throughout the lifespan of a captive quail population. This avian model has the particularity of being able to sustain a nearly constant egg laying profile for many weeks while under a photo-stimulatory photoperiod and no feed limitations (Satterlee and Marin, 2004). The study considers four experimental stages: i) juveniles and ii) adult quail; and then, iii) immediately after they have experienced an unpredictable chronic stress challenge (CS), and iv) 10 weeks later, during an advanced egg-laying stage. Thus, each evaluated stage provides distinct and complementary information. The first two stages or sampling points are associated with physiological changes linked to natural life processes, such as sexual maturation and growth (Garland et al., 2022). Using the second sampling point (adults before CS exposure) as a baseline, the third and fourth stages (adults after CS exposure and advanced egg-laying individuals) provide insights on the individuals’ ability to cope with external challenges in the short term (post-CS, third sampling point). The condition of challenged individuals at six-months of age (the last sampling point) can be understood both as an advanced egg-laying stage, supporting an elevated metabolic demand associated with the egg laying (Woodard and Abplanalp, 1971), and, possibly, as an indicator of medium- to long-term CS effects.

## Materials and Methods

### Animals and husbandry

We used Japanese quail (n=96, 48 males and 48 females) because they are recognized as an useful laboratory avian model due to their physiology, small size, short life cycle, and low maintenance cost (Huss et al., 2008). The quail were raised as described elsewhere (Nazar et al. 2012; Caliva et al. 2017). Animal care was provided in adherence to Institutional Animal Care and Use Committee guidelines.

Briefly, after hatch, quail were housed in enriched rearing boxes measuring 90 x 90 x 60 cm (length x width x height) in mixed groups of 40 individuals. A wire-mesh floor (1 cm grid) was raised 5 cm to allow excreta passage, and 50% of the total surface was covered with corrugated cardboard. Each box had a feeder arranged in the front, 16 automatic drinkers (“nipple” type), and a cover lid to prevent the escape of the birds and the loss of heat. The breeding temperature was lowered gradually and weekly from hatching to 4 weeks of age, starting from an initial temperature of 37°C until reaching 24°C (Nazar et al. 2018). The ambient relative humidity recorded ranged between 50 and 60%. The quail were kept in a daily cycle of 14 h of light (330-350 lx) and 10 h of dark (14:10 L:D) during the study, turning on the lights at 06:00 h. At 28 DA, quail were sexed based on breast plumage coloration and wing-banded for later identification. At 5 weeks of age, 48 male-female pairs were randomly housed in pedigree enclosures measuring 39 x 19 x 25 cm (length x width x height). The environmental enrichment program included corrugated cardboard and plastic grass on the floor, as well as rough plastic sheets and wooden boards, which were alternated weekly to minimize habituation and prevent disturbance from more frequent changes (Nazar and Marin 2011). A quail starter diet (24% crude protein; 2900 kcal metabolizable energy kg−1) was provided for the first 4 weeks of life, followed by a laying diet (21% protein, 2750 kcal ME kg−1). Water and food were supplied *ad libitum*.

### Study design

The study evaluated selected INE representative variables (previously associated with INPs, see below) by taking blood samples from each individual in four target stages (Fig. 2): 1) juvenile, pre-pubescent, quail (6 wks of age); 2) adult quail (15 wks of age), when birds evidenced full sexual maturity (all males producing foam from cloacal gland, and all females were at peak egg production (∼one egg per day), and immediately before an exposure to a chronic disruptive event (hereafter adults pre-stress; Table 1); 3) after chronic stress exposure (hereafter adults post-stress; 17 wks. of age); and 4) at 27 wks. of age, during an advanced egg-laying stage (six-month old), when decreases in peak egg production, fertility, and hatchability have already been shown (Woodard and Abplanalp, 1971).

**Fig. 2:**
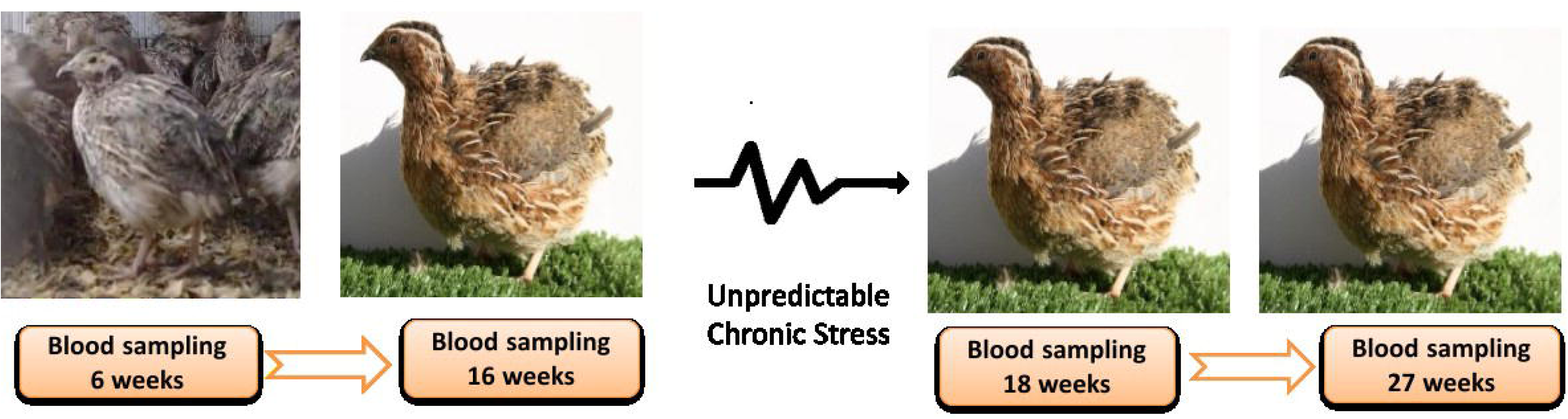
Summary of the experimental protocol followed in this research. INPs were assessed in a group of 96 quail during four experimental stages: 1) juveniles, 2) adults before an unpredictable chronic stress protocol, 3) adults after an unpredictable chronic stress protocol, 4) advanced egg-laying. Abbreviations: wk. (weeks), UCSE (Unpredictable Chronic Stress Exposure).

**Table 1:**
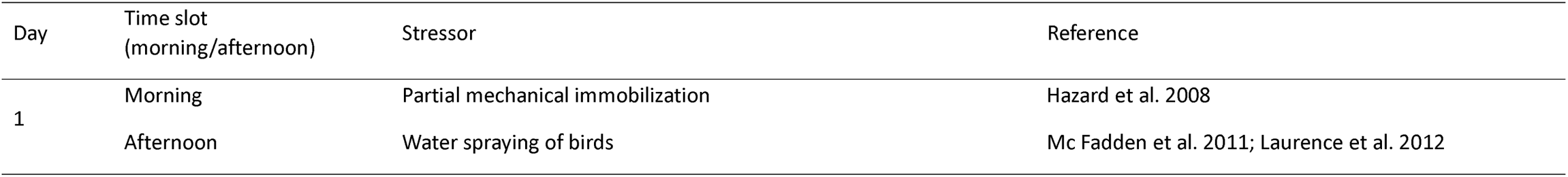
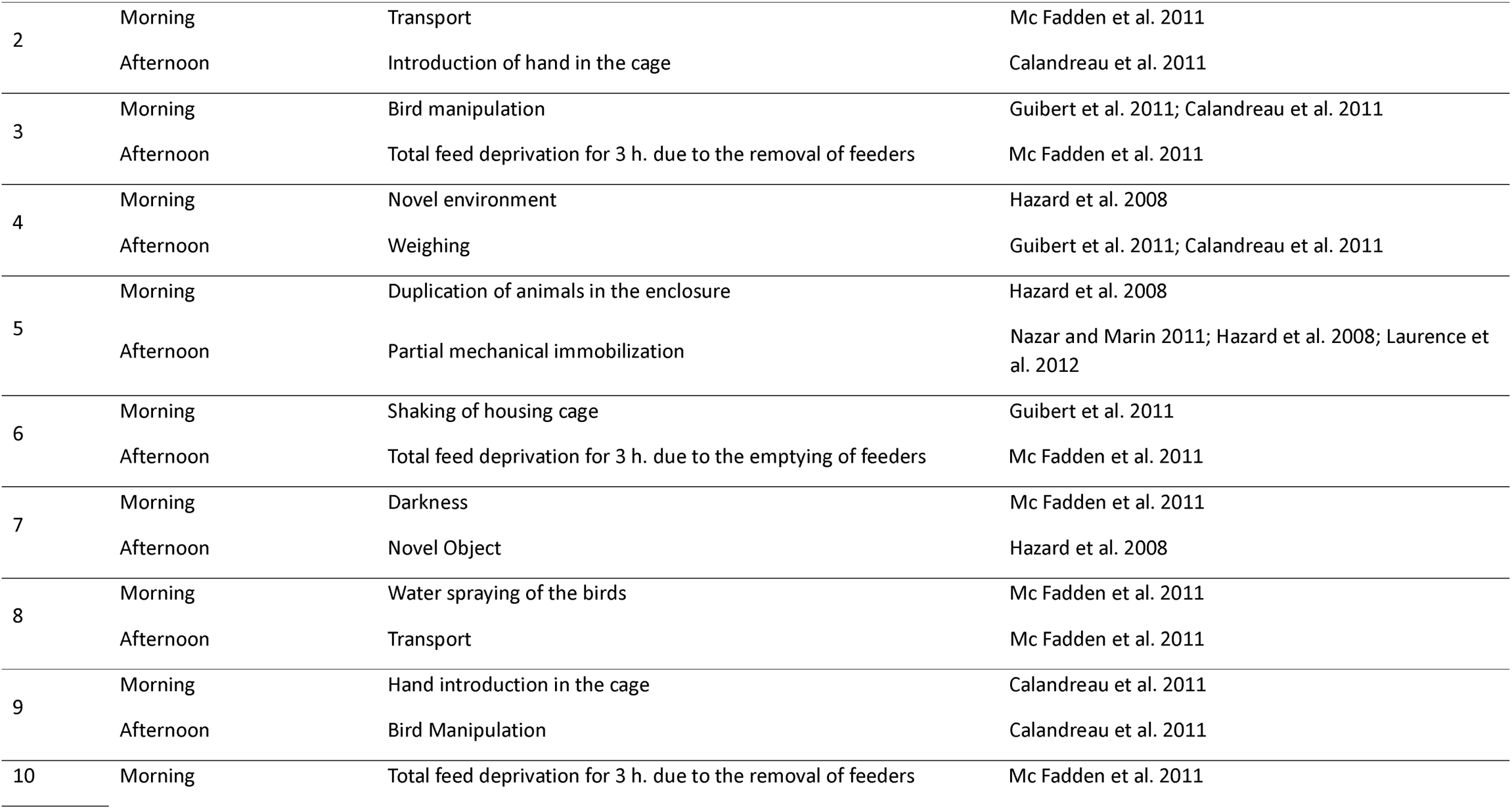
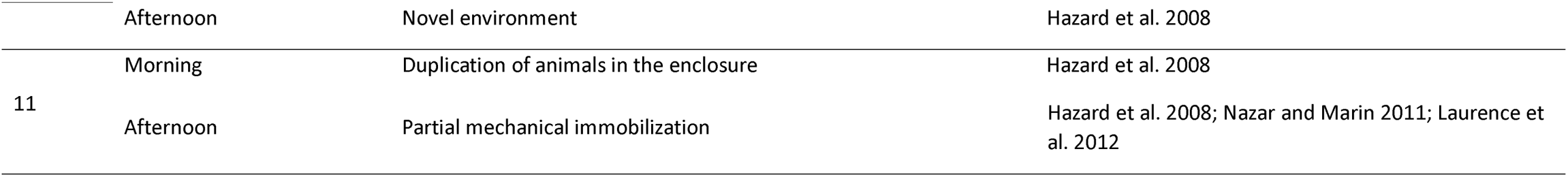
Unpredictable chronic stress exposure protocol developed in quail to determine its effect on the population distribution of INPs. Two different Stressors were applied each day, one during the morning hours (8:00 - 12:00 h) and the other in the afternoon hours (13:00 - 18:00 h). The exact time in which the stressor was applied varied from day to day to minimize predictability.

### Variables tested

The set of variables evaluated for characterizing INPs in every individual are briefly described below as fully descriptions are provided elsewhere (Nazar 2015; Nazar et al. 2015; Nazar et al. 2017).

#### Response to PHA-P injection

this is a measure of the birdś pro-inflammatory potential. Briefly, after measuring the wing web thickness with a caliper, a 0.05 ml intradermal injection of a 1 mg/ml solution of PHA-P in phosphate-buffered saline (PBS) was administered to the wing web area. The dermal swelling response was measured 24 h later as the percentage of increase in wing web thickness at the injection site. Measurements were recorded to the nearest 0.01 mm using a mechanical micrometer (Vinkler et al., 2010).

#### Antibody response to SRBC

This parameter was used to evaluate the induced humoral immune response. The antibody titers were assessed with a micro agglutination assay (Nazar and Marin 2011). A 20 μl sample of complement-inactivated plasma (heated to 56°C) was serially diluted in 20 μl of PBS (1:2, 1:4, 1:8, up to 1:512). Then, 20 μl of a 2% suspension of SRBC in PBS were added to all wells. Microplates were incubated at 40°C for 1 h, and hemagglutination of test plasma samples was compared to the blank (PBS only) and negative controls (wells with no SRBC suspension). Antibody titers were reported as the log_2_ of the highest dilution yielding significant agglutination. The results represent the average of two duplicates per animal.

#### Corticosterone and testosterone analyses

Plasma samples were extracted with diethylether (10 times the volume). The ether phase was transferred into a new vial, evaporated, and re-dissolved in assay buffer (Schöffmann et al., 2009). Aliquots (in duplicate) were analyzed using enzyme immunoassay (EIA) for corticosterone and testosterone. Details information regarding the EIAs, including antibody cross-reactivities, are given elsewhere (Palme and Möstl 1997; Auer et al. 2020).

#### Frequency of leukocyte subpopulation distribution (FLD)

Leukocyte subpopulations were quantified by flow cytometry. To perform this, 40 μl of total blood was stained with a fluorescent lipophilic dye (3, 3’-dyhexiloxacarbocyanine iodide; DiOC6, Molecular Probes) to obtain absolute counts of erythrocytes, lymphocytes, monocytes, thrombocytes, and granulocytes, as described elsewhere (Uchiyama et al., 2005). Then, the FLD number was calculated using the following formula: FLD = number of granulocytes/(number of lymphocytes + number of monocytes), serving as an indicator of the innate to acquired immunity ratio.

#### Cytokines

To determine cytokine mRNA expression levels, peripheral lymphocytes were isolated, and total RNA was extracted according to the manufacturer’s instructions, as previously described (Shini and Kaiser, 2009). The total RNA was reverse-transcribed, and the resulting cDNA was stored at −80°C until used for real-time PCR. For quantitative real-time PCR assays, specific primers for quail IL-1β, IL-4, IL-13, and IFN-γ genes were used as previously reported (Superina et al., 2009). β-actin was used as the reference housekeeping gene. Real-time PCR was performed with a StepOnePlus Detection System (Thermo Fisher Scientific). Each analysis was performed in triplicate for every bird, and the results represent the average of these replicates. The level of expression of each target gene was calculated using the formula: Gene Level = 2-(Target Gene Ct – β-Actin Ct). The value obtained was used to compare the level of expression of each molecule (Nazar et al. 2015).

### Statistical analysis

To assign specific INPs to each individual, we conducted a cluster analysis to explore the degree of similarity among the birds based on the analyzed variables, following Nazar *et al.* (2017). Specifically, to identify the most sensible partition of the birds, described by the best orthogonal linear combinations of factors using the least-squares criterion, a K-means analysis was performed using Infostat software (version 2020, https://www.infostat.com.ar/). The analysis included the following 9 variables: plasma corticosterone, testosterone, IFN-γ and IL-4, IL-1β, IL-13, skin-swelling response to PHA-P, antibody response against SRBC, and FLD ratio. Based on the theoretical framework previously exposed, the clustering was performed with a K-value of 3 (2 subgroups with representatives of potential extreme INPs and 1 intermediate subgroup). Euclidean metrics were used for computing the distance between points and cluster centers. To graphically represent the clustering results in a multivariate manner, a Principal Component Analysis (PCA) was performed using the clustering variables as explanatory variables (Crawley 2007; Nazar et al. 2017; R Core Team 2021).

We evaluated the transition probability from one INP to the others during the four experimental stages using discrete-time Markov chains (DTMC; Spedicato and Signorelli, 2014) using R v.3.6.3. This method is well-suited for analyzing changes in individuals over successive time points. Transition matrices were created for each pair of stages: Specifically, we created: (i) a matrix comparing juveniles to adults, (ii) a matrix comparing pre-stress to post-stress adults, and (iii) a matrix comparing post-stress adults to individuals in the advanced egg-laying stage (Bishop et al., 1974). Transition matrices estimate the probability of an individual changing its phenotype. For example, if 16 individuals show a Fischer-like phenotype at the juvenile stage, we aimed to determine how many of them remained Fischer-like upon reaching adulthood and how many transitioned to Lewis-like or the intermediate phenotype. We also evaluated how many individuals retained the same phenotype across stages. To estimate probabilities, we divided the number of individuals that changed phenotypes by the total of individuals with that phenotype in the previous state (Spedicato and Signorelli, 2014). In 27 cases along the four experimental stages, the INP could not be established at all four stages, which may cause a slight variation in the total number of individuals per transition. However, for all three transitions, the sample size always exceeded 70 individuals, making the results representative of INP configurations in the species and their changes throughout the experiment.

## Results

K-means analysis grouped the birds in 3 main dissimilar clusters across the four experimental stages. The three groups identified by the cluster analysis are represented in the PCA plots (Fig.3), using biplots as a graphical strategy to represent the observed phenomena. The three groups remained differentiated throughout four experiment stages, although changes in frequency were also observed. In the first two stages (juveniles and adults pre-stress), the intermediate phenotype had a higher number of individuals compared to the extreme phenotypes, i.e., Fischer-like and Lewis-like, as shown in the PCA plots (Fig. 3). The next stages (adults post-stress and six-month old individuals) showed a more even INP distribution (Fig 3).

**Fig. 3:**
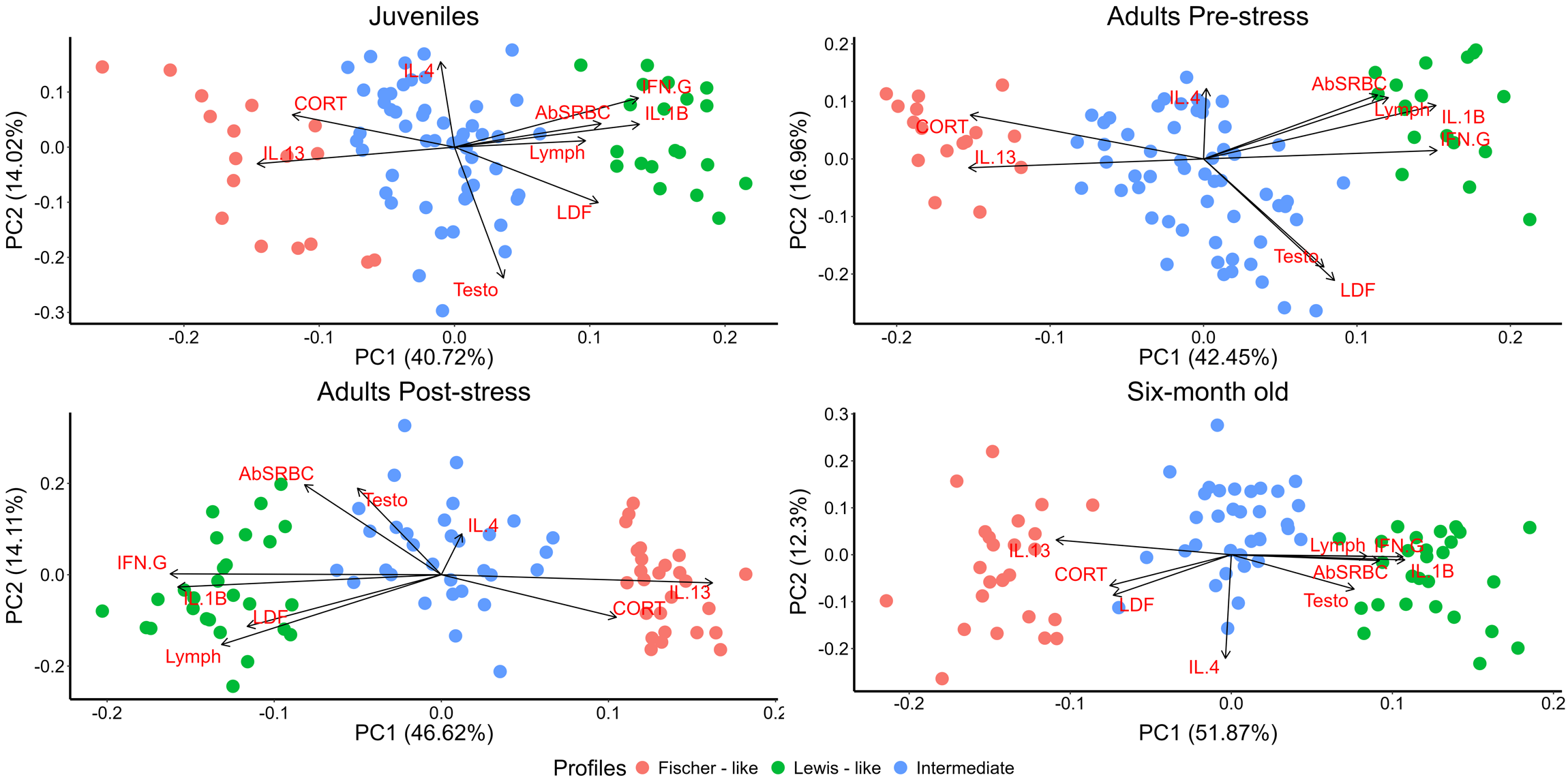
Principal component analysis plots exploring data variability and clustering procedure. Each dot represents one bird and each variable is represented by an arrow. Each plot represents the INP during one stage of the experiment.

The frequency of the INPs in the studied population showed that the intermediate phenotype, characterized by moderate INE responses, was predominant during the first two experimental stages (Fig. 4). However, after stress exposure, the frequency of the extreme phenotypes, Lewis-like and Fischer-like, significantly increased (Table 2; p-value = 0.04 for Lewis-like and p-value = 0.02 for Fischer-like). Simultaneously, there was a marked reduction in the proportion of individuals with intermediate phenotype during this stage (p-value < 0.01). In the advanced six-month old individuals (advanced egg laying stage), the frequencies of the extreme phenotypes remained stable compared to the previous (post-stress) stage. Table 2 shows the results of the proportion test comparing transitions among phenotypes. A significant increase in the proportion of extreme phenotypes, alongside a reduction of the intermediate phenotype, was observed only in the transition from adults to post-stress adults (highlighted in bold). No main differences were found in any of the other transitions.

**Fig. 4:**
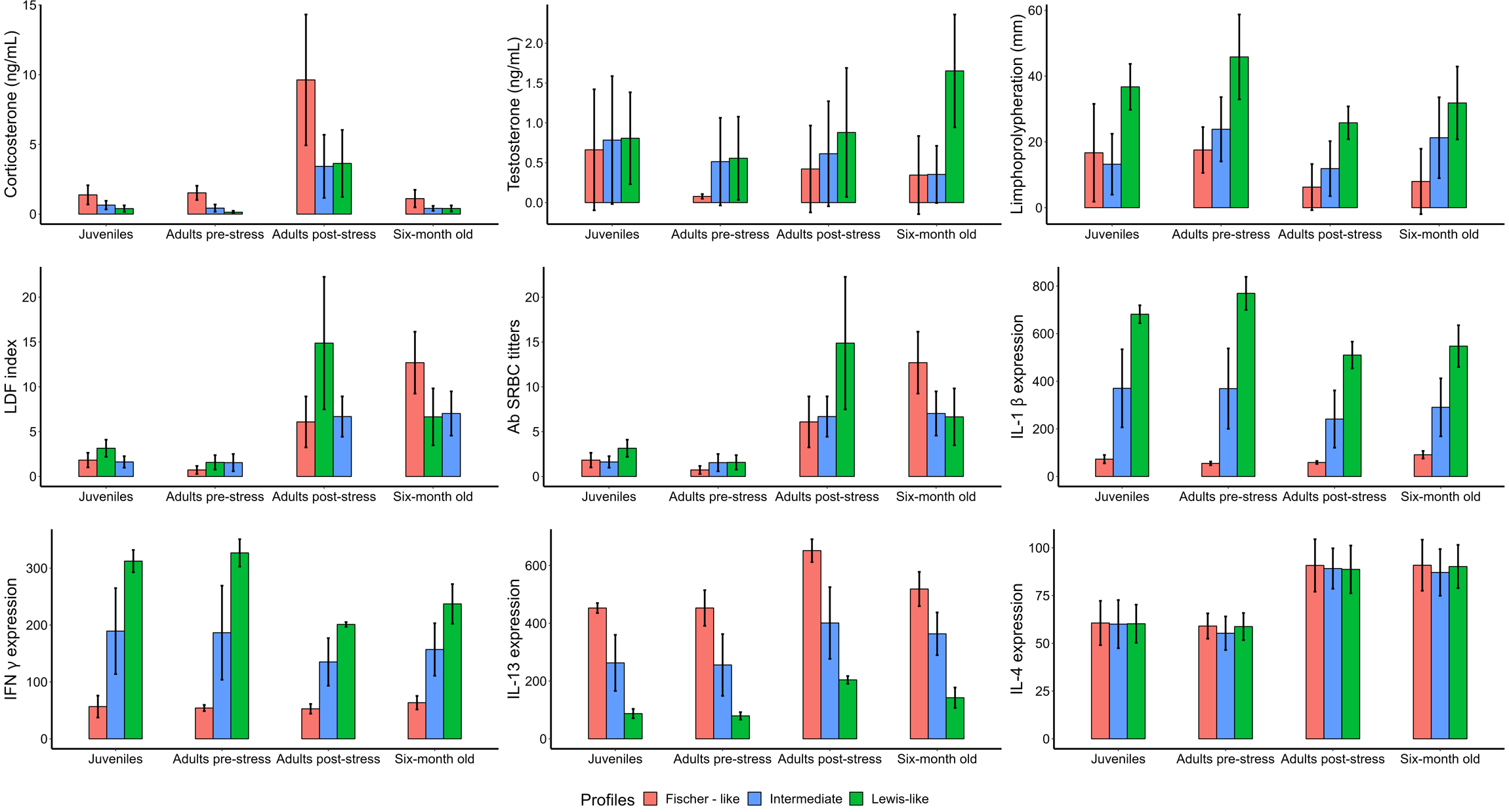
Linear discriminant analysis plots. Each plot represents the INP during one stage of the experiment. The variables Limph, Testosterone, AbSRBC, IFN-G and IL.4 showed similar values along canonical axis 1 and 2, resulting in their overlap near the center of the graph. In the “adult post-stress” plot, corticosterone also showed a similar displacement along the canonical axes.

**Table 2:**
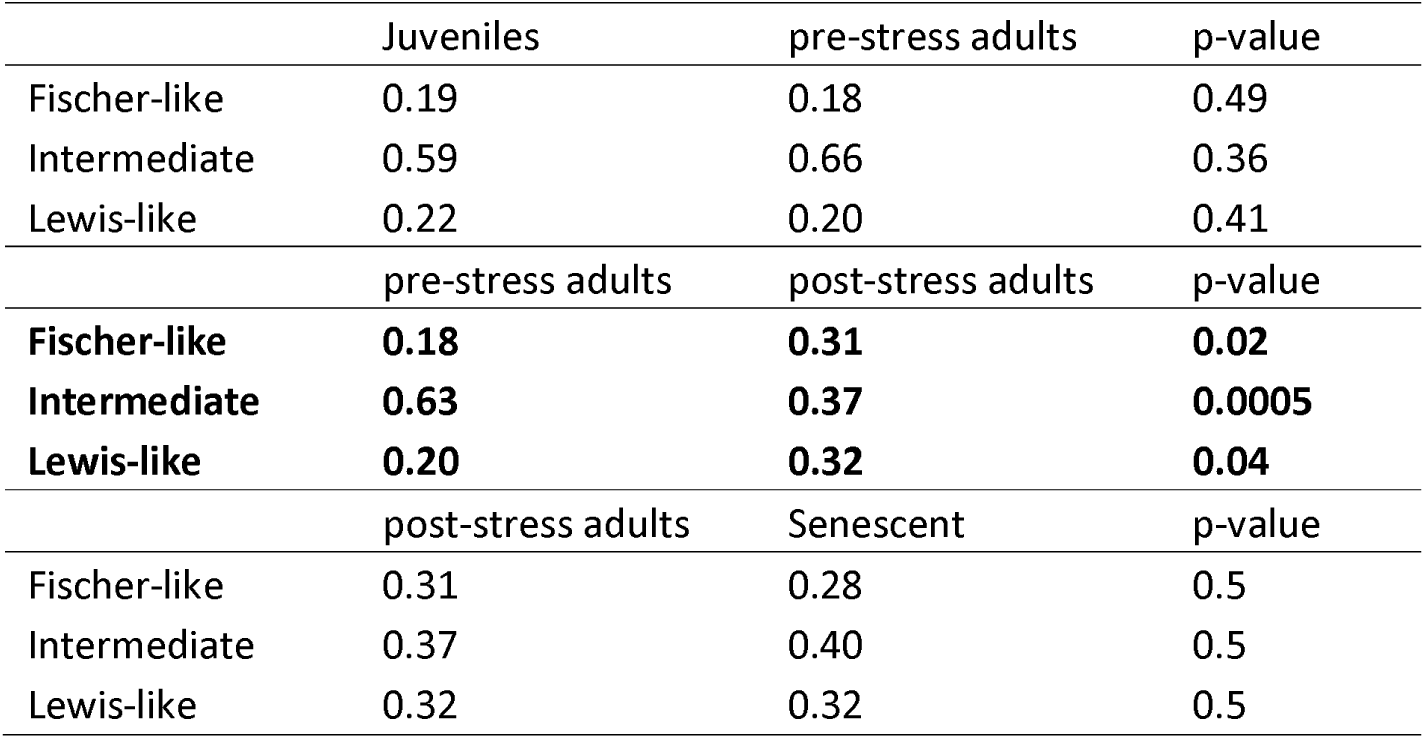
Proportion analysis to assess differences among phenotypes in the three ontogenetic transitions studied.

The proportion analysis comparing the phenotypes among sexes within the same experimental stage (Table 3) indicated that following the stress protocol, the number of individuals displaying Fischer and Lewis like phenotypes increased, and this shift remained consistent through to the advanced egg-laying stage (Fig. 3). Additionally, the proportion of females showing a Fischer-like phenotype was significatively higher than males with this INP in adults pre-stress (p-value < 0.01, Table 2), in adults post-stress (p-value < 0.01), and in advanced egg-laying quail (p-value <0.01). On the contrary, the proportion of males presenting a Lewis-like INP was significantly higher than the proportion of females in the adults’ pre-stress (p-value < 0.01), and in the advanced egg-laying experimental stages (p-value < 0.01). In addition, the frequency of males with an intermediate phenotype was significantly higher than the proportion of females in the adults’ post-stress experimental stage (p-value < 0.01). Only during the juvenile assessment, a trend for females showing a reduced proportion of Lewis-like phenotype compared to males was found (Table 2).

**Table 3:**
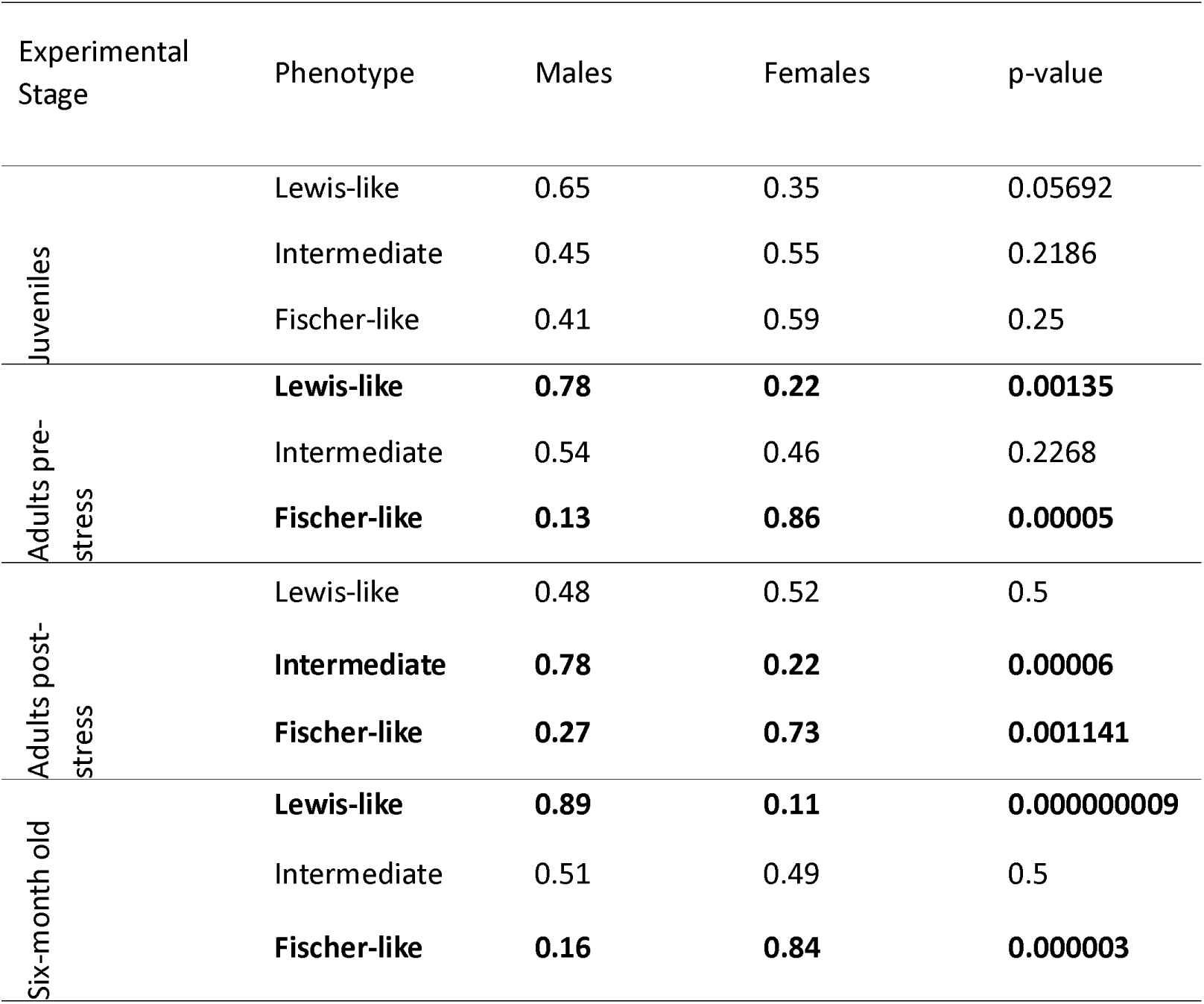
Proportion analysis comparing differences among sexes during the four experimental stages: juveniles, adults pre-stress, adults post-stress and senescent. Significative differences are indicated in bold.

The probability of an individual changing its phenotype between experimental stages, or remaining in the same phenotype, was evaluated using a DTMC analysis (Table 4). During the transition from juveniles to adults, the highest probabilities of change were observed for individuals shifting from Lewis-like or Fischer-like phenotypes to the intermediate phenotype (0.55 and 0.75, respectively). In contrast, most individuals classified as intermediate remained in that phenotype (0.64; Table 3; Fig. 5). After the stress protocol, the probability of individuals shifting from the Lewis-like (0.36) or Fischer-like (0.40) phenotypes to the intermediate phenotype decreased (Table 3; Fig. 5). However, the probability of individuals shifting from the intermediate phenotype to either Lewis-like (0.41) or Fischer-like (0.29) increased. Additionally, the probability of Lewis-like individuals changing their phenotype to Fischer-like, and vice versa (0.27 and 0.33, respectively), also increased during this transition. Finally, in the transition from post-stress adults to advanced egg-laying individuals (Table 4), the probability of individuals transitioning from the intermediate phenotype to Lewis-like remained relatively high (0.44). The probability of changing between Lewis-like and Fischer-like phenotypes (0.20 to both of them) decreased compared to the pre-stress to post-stress adult transition.

**Table 4:**
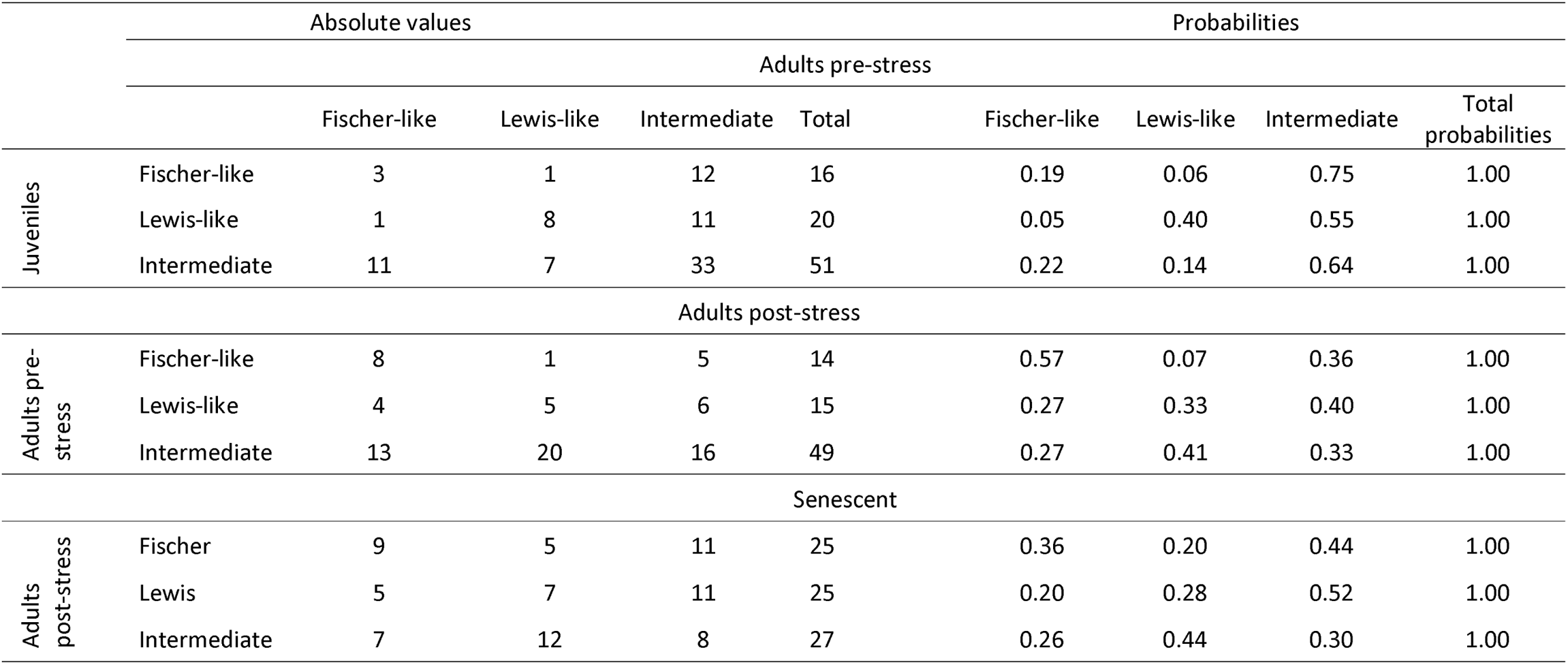
Transition matrices estimating the probability of an individual changing its phenotype from one experimental stage to the other or the probability of remaining in their original phenotype. On the left are represented the number (in absolute values) of individuals that belong to a determined INP and remained there in the next stage or had a different INP during the next stage On the right is represented the transitions probabilities matrix created with data on the left. Transitions represented: from juveniles to adults pre-stress, from adults pre-stress to adults post-stress, and adults post-stress to senescence. Note that for DTMC analysis probabilities were calculated considering the partial totals for each phenotype in each stage.

**Fig. 5:** Phenotype frequency distribution during the four stages of the experiment. Each plot represents the INPs frequency during one stage of the experiment.

**Fig. 6:** Plot of transition probabilities. Arrows indicate changes from one phenotype to the other or the permanence in that INP. Transition probabilities are indicated on the lines.

## Discussion

This case study assesses changes in INE variables that lead to the configuration of INPs throughout different life events. These events include sexual development (represented in our design by juveniles), coping with adult life demands (represented by adults pre-stress), experiencing unpredictable chronic stress events during adulthood (adults post-stress), and, finally, reaching an advanced egg laying stage more than two months after CS exposure (six-month-old individuals). All of these situations can occur during an individual’s lifespan and appear to influence INP proportions, their distribution at the population level, and their probability of change. It is crucial to emphasize that herein INPs are not considered fixed entities here but rather represent the array of INE responses that an individual can deploy at a given point in its ontogeny.

It has been proposed that the stress response is influenced by both the life history traits of the species (Schoenle et al., 2021) and the potential response of different tissues (Lattin et al., 2013; Scanes, 2022). Furthermore, INPs have been suggested as an integrative factor to consider in this framework (Elenkov et al., 2008; E. M. Sternberg et al., 1989; E M Sternberg et al., 1989; Wei et al., 2003; Zelazowski et al., 1992). Fischer-like individuals are more associated with anti-inflammatory and antibody-mediated immune responses than Lewis-like individuals (Nazar et al. 2017; Nazar et al. 2015; Mosmann et al. 1986). Therefore, individuals expressing either of these two extreme phenotypes likely possess different abilities to cope with environmental conditions, showing distinct sensitivities to challenges such as stressors (Colditz et al., 2016). Changes in INP configuration from one life stage to another may then suggest a shift in the individual’s coping abilities (Zelazowski et al. 1992; Wei, Listwak, and Sternberg 2003; Nazar et al. 2015; Nazar et al. 2017). Theoretically, if an individual with an intermediate INP prior to stress shifts to a Lewis-like phenotype following chronic stress, its ability to develop robust inflammatory responses increases, enhancing its capacity to deal with certain types of pathogens (Blom and Ottaviani, 2017; Elenkov et al., 2008; Lee, 2006; Nazar et al., 2017; Wei et al., 2003). Conversely, if an individual acquires a Fischer-like configuration, its response to infections may differ substantially from that of a Lewis-like individual (Wei, Listwak, and Sternberg 2003; Elenkov et al. 2008; Nazar et al. 2017). These examples illustrate some of the observed shifts, but it is crucial to understand that any INP shift results from a change in the integration of the components of the INE system and thus reflects a shift in the potential responses an individual can deploy.

In the juvenile and adult pre-CS stages, the frequency of extreme phenotypes was low, while the frequency of the intermediate phenotype was notoriously higher (Table 3). However, no significant increase in phenotype frequencies was observed during the transition from juveniles to adults (Table 1). In contrast, after CS, the newly configured population displayed a significant increase in extreme phenotypes and a corresponding decline in intermediate ones (Table 1). Consequently, the frequency of extremes INP became almost equal to the intermediate INP. It is important to recall that the stress protocol included not only various stressors but also a high degree of unpredictability. Therefore, the shift toward an increased representation of extreme INPs could be seen as an adaptive strategy to manage a wider range of environmental and immune challenges, whereas a population dominated by a single INP might have limited adaptability (Blom and Ottaviani, 2017; Forsman, 2015; Nazar et al., 2015). Remarkably, this new distribution remained stable through the advanced egg-laying stage (Table 3), with no significant changes in the proportion of individuals belonging to any particular INP. While this study design makes it difficult to differentiate the effects of stress from aging, it is likely that a combination of both factors influenced INP configuration in six-month-old individuals. These findings suggest that the INP changes, and therefore the populatiońs ability to cope with future life events, may be related to long-lasting carry-over effects, as previously proposed in the literature (Senner, Conklin, and Piersma 2015; Nazar et al. 2017; Moore and Martin 2019).

The proportion of females expressing the Fischer-like phenotype was significantly higher than the proportion of males in the last three sampling points (adult pre-CS, post-CS, and six-months stages). In contrast, males exhibited a higher frequency of the Lewis-like phenotype during those stages. These findings provide insight into how both sexes cope with challenges, indicating a polarization in INP expression, with males more frequently expressing a Th1 response than females, suggesting greater preparedness for intracellular infections (Kogut, 2022; Lee, 2006). This polarization may be related to differing energetic demands of reproduction in males and females previously mentioned: males tend to invest more energy in mate attraction, which may reduce their investment in immune responses to parasites (Lee, 2006; Valdebenito et al., 2021). Furthermore, studies in broilers and zebra finches have respectively shown that males exhibit a higher corticosterone response to stress (Marin et al., 2002) and a lower survival rate after a stress induction compared to females (Jimeno et al., 2018). Given that Lewis-like individuals have lower basal corticosterone than Fischer-like or intermediate individuals, the increased prevalence of males with this phenotype might be advantageous. These males may respond differently to future challenges, potentially lowering the demands associated with their stress responses while enhancing other traits such as fitness or survival. This proposal requires further investigation, and the relation between INPs and sex will be the focus of specific future research.

From the individual’s perspective, most individuals exhibited changes in their INP configurations during the experiment, even in the absence of environmental challenges (Fig. 5). Phenotypic changes over an individual’s lifetime can progress linearly, particularly during juvenile stages, affecting subsequent developmental steps. In contrast, during adulthood, these changes may occur in a more flexible and adaptive manner in response to environmental conditions (Jacobs and Wingfield, 2000; Senner et al., 2015). The ability to vary INP in response to environmental factors could represent a physiological advantage in certain cases. However, an important question arises: do these changes increase the burden on the individual? This likely depends on the complex interaction between ontogenetic demands, the individual’s energy budget, and environmental challenges (Garland et al., 2022; Shankar et al., 2019; Walsberg, 2003). Although all individuals in our experiment had the same access to nutritional resources, this does not mean that their energy utilization was identical. As previously noted, some individuals may have a greater capacity to allocate energy across various physiological processes, enabling them to intake, digest, and utilize greater amounts of energy than their conspecifics (Buehler et al., 2010; Halsey et al., 2019). Ontogenetic variations, such as sexual maturation and reproduction, may influence other physiological systems, including the INE matrix, leading to shifts in INPs throughout life (Johnston et al., 2012; Segner et al., 2017). These endogenous disruptors can influence the individual without necessarily altering the overall population structure, which, as shown here, may remain balanced.

INP changes may reflect a shift in INE responses and potential, which could imply a trade-off in energy allocation (Garland et al., 2022). The extent and possibilities of this trade-off are limited by how efficiently individuals can utilize energy at any given moment (Buehler et al., 2010; Hasselquist and Nilsson, 2012; Yang et al., 2021). In this case study, some individuals may exhibit the effects of CS demands immediately after the experiment, while others could display carry-over effects in response to the challenges (Senner et al., 2015). However, since all transition probabilities seem to increase after the stress protocol, individuals may become more flexible after experiencing stress. Adapting their INP might be more feasible after CS, potentially providing an advantage for coping with environmental complexity and demands (Pirotta et al., 2022).

In recent years, there has been growing interest in how individuals will cope with environmental changes, such as weather variations linked to climate change and its broader effects (Burggren, 2018; Cunningham et al., 2021). Increased plasticity in the INE system may provide a significant advantage by allowing individuals to adapt to diverse and challenging situations (Cockrem, 2013). Moreover, as temperature fluctuations may affect pathogens diversity and richness, greater INP flexibility, as shown here, could enhance an individuaĺs ability to manage such threats (Sergio et al., 2018). Understanding how challenges are integrated at the individual level can provide valuable insights, while the collective responses of individuals within a population, as seen in our study, offer a new perspective on population dynamics and adaptive potential.

In summary, this case study provides new insights into INP changes at multiple scale of analysis (individual or population levels) showing that INP expression may be variable according to internal and external changes. Even in a highly selected species like quail, which exhibit an impressive capacity to sustain an egg-laying rates close to 95% of physiological maximum, individuals retain the capacity to adjust their physiological responses throughout their lifespan, adapting to internal changes such as sexual maturation or aging. However, when viewed at the population level, these gradual ontogenetic shifts appear less pronounced, as the overall INP distribution remains relatively stable over time. In contrast, exposure to chronic stress acts as a powerful external force, reshaping the population structure by increasing INP variability and altering the balance between phenotypes. This suggest that wile individuals may show flexibility in their physiological responses, large-scale population shifts are more likely to occur in response to external pressures rather than internal developmental processes. Understanding these dynamics is crucial for predicting how populations may adapt to changing environmental conditions and for developing management strategies that account for both individual plasticity and collective resilience.

## Acknowledgments

This research was done with funding from the National Agency of Scientific and Technological Research (PICT: 2016-1979). We would like to thank M. Julia Ortiz, Pablo Prokopiuk and Dario Arbelo for their technical assistance during the development of the experiment, and Edith Klobetz-Rassam for EIA analyses.

## Authors Contributions

Experimental design: F.N.N, R.H.M., S.G.C., A.M. Laboratory Analysis: M.F.P, V.I.C; R.P; FNN. Statistical Analysis and figures preparation: A.M; Original draft writing: A.M., F.N.N. Manuscript writing and revision: R.H.M., S.G.C; R.P. V.I.C, M.F.P. All authors have reviewed and approved the final manuscript.

## Data Availability

Data is available at https://data.mendeley.com/datasets/vs48x3x9f2/1. Contact

## Ethical statement

The experiment procedures were approved by the ethical committee at Instituto de Investigaciones Biológicas y Tecnológicas (IIByT) in compliance with the legislation regarding the use of animals for experimental and other scientific purposes (ACTA CICUAL N° 27, 09/04/2015).

